# Stereochemical identity of lipid nanoparticles modulates protein expression via internal lipid organization

**DOI:** 10.64898/2026.06.05.730351

**Authors:** Dennis Aschmann, Renzo A. Knol, Wessel Verbeet, Sven Wijngaarden, Oscar Escalona-Rayo, Rafael V.M. Freire, Karl Bertram, Indira Tekkali, Georg Bunzel, Bauke L. Fontein, Aiswarya Dharan, Inès Pfister, Yun Zhang, Tom Keijer, Joost N. H. Reek, Ilja K. Voets, Bram Slütter, Alexander Kros

## Abstract

Stereochemistry plays a crucial role in how molecules interact with complex physiological environments, affecting pharmacokinetics, pharmacodynamics, efficacy, and toxicity. Although these effects are well studied for small-molecular drugs, they are largely overlooked for supramolecular assemblies used in drug delivery. Even for lipid nanoparticles (LNPs)—the most advanced RNA delivery platform—stereochemical effects are rarely investigated and, when considered, are typically limited to the ionizable lipid rather than the overall stereochemical identity of the LNP.

Here we separate the ionizable lipid cKK-E12 into its two stereoisomers (*trans*: *R,S*/*S,R*; *cis*: *R,R*/*S,S*), which are normally used as a mixture. LNPs containing the *cis* isomer exhibit improved physicochemical properties, stability, and protein expression. By systematically varying the stereochemistry of the ionizable lipid, phospholipid, and cholesterol, we reveal stereochemistry-dependent differences in uptake and protein expression across six cell lines and *in vivo* in zebrafish embryos and mice. AI-assisted cryo-TEM analysis and SAXS link enhanced protein expression to structural differences, demonstrating control over internal lipid phases (lamellar and inverse hexagonal), influencing sample uniformity, and identifying stereochemical identity as a key determinant of functional RNA delivery.

## Introduction

The impact of stereochemistry is well established in the field of small-molecule drugs. Since the tragic example of Thalidomide (Contergan) in the late 1950s—administered to pregnant women as a sedative but causing severe birth defects due to racemization—the importance of stereochemical purity in pharmaceuticals has become widely recognized.^1,2^ It is now well understood that stereochemistry can influence pharmacokinetics, pharmacodynamics, potency, efficacy, as well as the overall safety profile of therapeutic agents.^3–7^

In contrast, much less is known about the role of stereochemistry in supramolecular assemblies used for drug delivery. Remarkably, even for the most advanced RNA delivery platform—lipid nanoparticles (LNPs), which became widely used as COVID-19 mRNA vaccines—stereochemical effects remain largely overlooked and are only briefly addressed in a few studies. This is surprising since LNPs consist of four lipid components (an ionizable lipid, cholesterol, a phospholipid, and a PEGylated lipid), several of which contain stereocenters. To date, the limited observations mainly attribute differences in protein expression to the ionizable lipid.^8–11^ For example, C12-200 has been synthesized with all hydroxyl groups in either the *S* or *R* configuration.^11^ Another study reported artificial lipids incorporating an amino acid to introduce chirality.^9^ In addition, batch-to-batch variations of cKK-E12 have been reported, while the molecular origins of these differences have not been systematically investigated.^10^ However, the potential contribution of the helper lipids remains largely unexplored, and the mechanisms by which stereochemistry influences expression levels are still unclear.

Here, we introduce the concept of stereochemical identity of LNPs, which captures the combined stereochemistry of all lipid components. We first demonstrate that cKK-E12 exhibits stereochemical variation arising from racemization at the diketopiperazine core. Using liquid chromatography, we separate cKK-E12 into two fractions containing either the *trans* isomers (*meso*: *R,S*/*S,R*) or the *cis* isomers (*R,R*/*S,S*). We then investigate how these isomers affect the performance and stability of the resulting LNPs. Finally, by replacing cholesterol (Chol) and 1,2-distearoyl-*sn*-glycero-3-phosphocholine (DSPC) with their complementary stereoisomers, we systematically examined how the stereochemical identity of LNPs influences physicochemical properties, protein expression, internal structure, and characteristic distance. Additionally, we investigated the effect of replacing DSPC with 1,2-dioleoyl-sn-glycero-3-phosphoethanolamine (DOPE), a previously used strategy to enhance protein expression through inverse hexagonal phase formation, increasing fusogenicity. ^12–16^

## Results

### Resolving the stereochemical composition of cKK-E12

To investigate the effect of stereochemistry on LNPs, we analyzed the ionizable lipid cKK-E12 by HPLC to determine whether its stereoisomers (diastereomers) could be separated, as the lipid contains six stereocenters (Fig. 1a). The chromatogram (Fig. 1e) revealed two distinct peaks with an intensity distribution of 60% (Peak 1) and 40% (Peak 2). Both peaks exhibited identical masses and nearly identical NMR spectra, indicating that they correspond to different stereoisomers (Figs. S1-S2, S11-S12). Although the four hydroxyl stereocenters could in principle generate up to 16 stereoisomers (2⁴), only two peaks were observed, suggesting that stereochemical variations at the DKP core dominate the resulting molecular structure. This suggests that stereochemical information at the diketopiperazine is lost during the final synthetic step due to the high temperature and strong basic conditions^17^, resulting in racemization of the stereochemically predefined diketopiperazine center and the formation of *cis* and *trans* isomers corresponding to the two observed peaks.

**Fig. 1.**
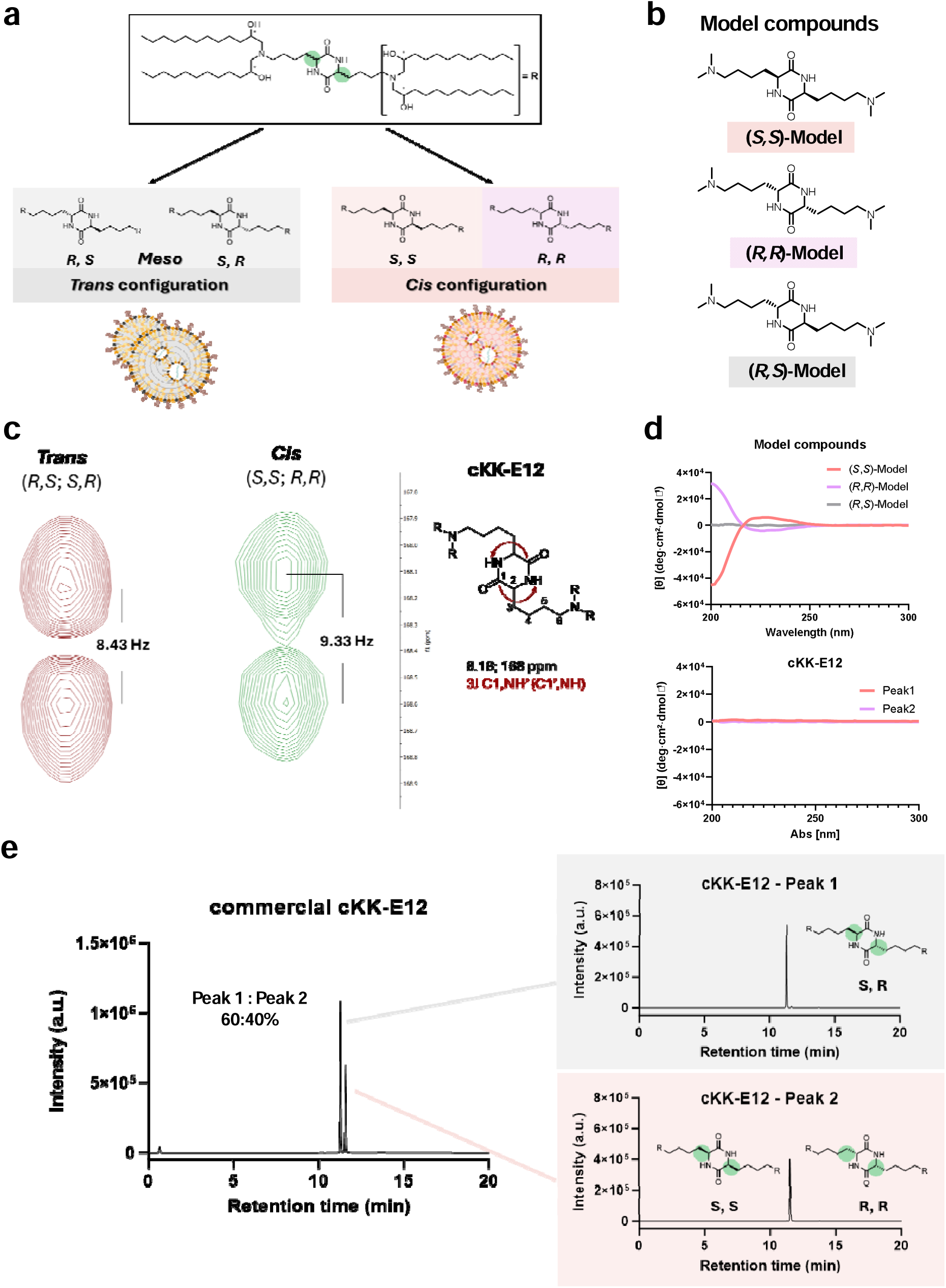
Deconvoluting the stereochemistry of different cKK-E12 fractions. **a**, Illustration of trans (*meso*) and *cis* configuration, leading to different morphologies and internal structures**. b**, Model compounds of cKK-E12 isomers. **c**, J-HMBC spectra of cKK-E12 Peak1 and Peak2 showing the ^3^*J*_CH_ coupling in DMSO-d6**. d**, CD spectra of model compounds and cKK-E12 Peaks**. e**, LC chromatogram of cKK-E12 mixture (commercial), allocating the stereochemistry to the peaks, detected by ELSD.

To test this hypothesis, we synthesized model compounds with the same core structure as cKK-E12 but lacking the lipid chains, representing either the *cis* (*R,R* and *S,S*) or the *trans* (*meso*, *R,S*) configuration. Circular dichroism (CD) spectra showed that the *R,R* and *S,S* models display opposite Cotton effects at ∼225 nm (*S,S* positive; *R,R* negative), whereas the *R,S* (*meso*) model exhibited no signal. Both cKK-E12 Peak 1 and Peak 2 also showed no CD signal, suggesting that one peak corresponds to the *meso* compound and the other to a racemic mixture of the *R,R* and *S,S* isomers, resulting in cancellation of the CD signal. To assign the peaks to the *trans* and *cis* isomers, we performed *J*-HMBC NMR experiments. These revealed ^3^*J*_CH_ coupling constants between the carbonyl carbon (C=O) and the amide NH proton of 8.43 Hz for the *trans* isomer and 9.33 Hz for the *cis* isomer (Fig. 1c). These values closely match those obtained for the model compounds, with 8.41 Hz for the *trans* model and 9.28 Hz for the averaged *R,R* and *S,S cis* models (Fig. S13).

Given that the *cis* and *trans* isomers of cKK-E12 are expected to influence lipid packing within LNPs (Fig. 1a), we subsequently formulated LNPs with varying isomeric ratios.

### *cis*-LNPs show superior encapsulation efficiency and stability to *trans*-LNPs

After successfully separating the cKK-E12 isomer variants, *cis*-cKK-E12 (racemic mixture of *R,R*-and *S,S*-cKK-E12) and *trans*-cKK-E12 (*meso* compound; *R,S*-cKK-E12), isomer-pure enhanced green fluorescent protein (EGFP)-mRNA-LNPs containing only one isomer variant (hereafter named *cis*-LNPs or *trans*-LNPs), as well as EGFP-mRNA-LNPs with defined ratios of the *cis* and *trans* isomers, including 25%, 50%, or 75% of either isomer, were prepared. These formulations contained DSPC, Chol, DMG-PEG2K, and DiD as a lipophilic dye at a fixed molar ratio of 50:10:37.3:2.5:0.2 (cKK-E12:DSPC:Chol:DMG-PEG2K:DiD). Additionally, a formulation containing the commercial cKK-E12 isomer mix served as a reference (Fig. 2a,b). The resulting LNPs were characterized by dynamic light scattering (DLS) to determine particle size, polydispersity index (PDI), and zeta potential. Interestingly, cKK-E12 isomer identity had a profound effect on these characteristics, with *cis*-cKK-E12 showing a smaller particle size and PDI, which increased with increasing *trans*-cKK-E12 content (Fig. 2c). Zeta potential showed negligible deviations between the groups. However, isomer mixtures generally showed slightly more negative zeta potentials than isomer-pure LNPs (Fig. 2d). Importantly, encapsulation efficiency was highest for *cis-*LNPs and decreased with increasing *trans*-cKK-E12 content (Fig. 2e).

**Fig. 2.**
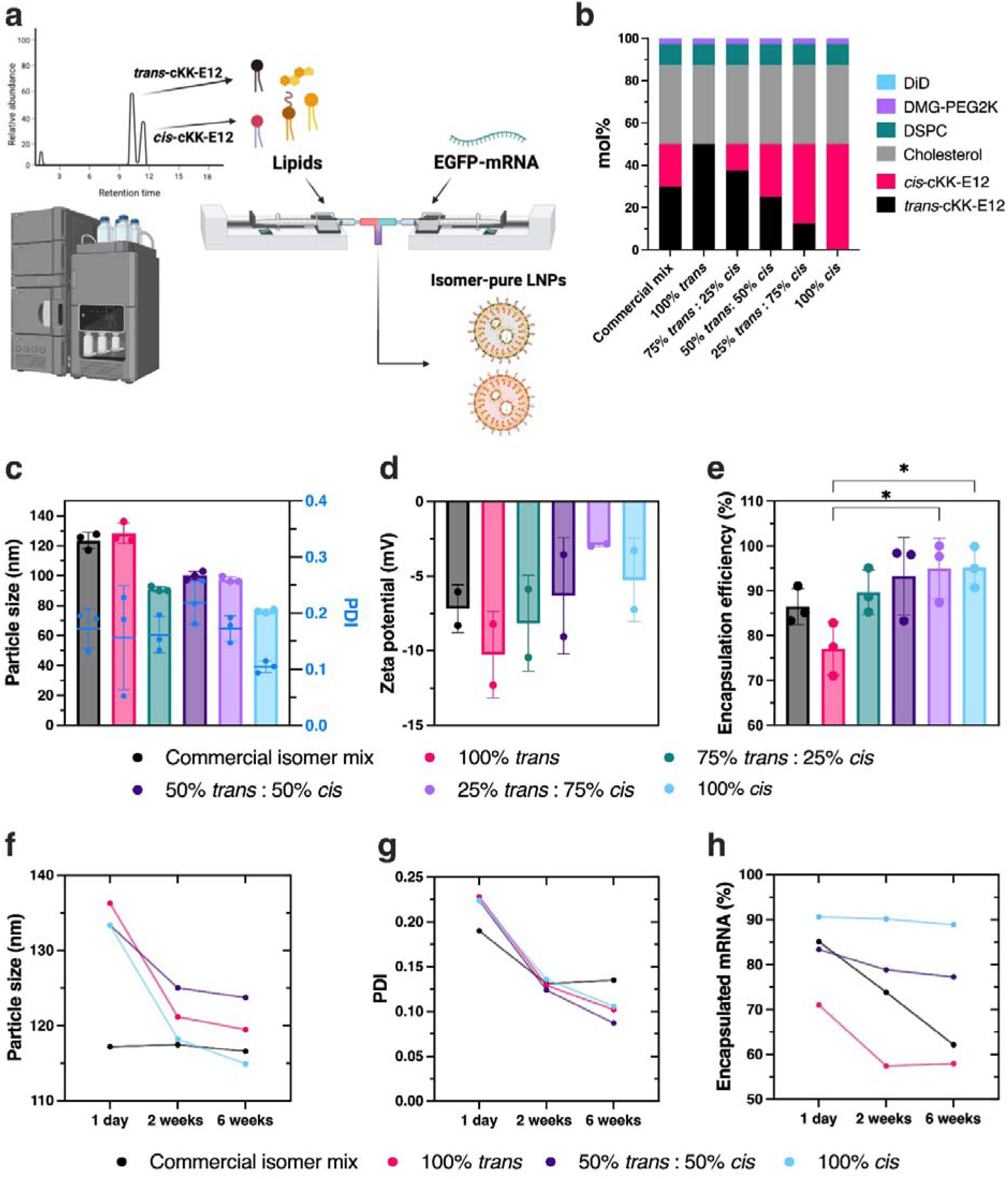
Manufacturing and physicochemical characterization of isomer-pure and isomer-mixed cKK-E12 lipid nanoparticles. **a**, Schematic representation of the LNP manufacturing procedure. **b**, Lipid content of formulations containing different cKK-E12 isomer variants. **c–e**, Particle size, polydispersity index (PDIs) (**c**), zeta potential (**d**), and encapsulation efficiency (**e**) of the LNP formulations. **f–h**, Physicochemical stability over 6 weeks of storage at 4°C: particle size (**f**), PDI (**g**), and encapsulation efficiency (**h**). Data is presented as group mean (n=3) ± SD and statistically compared by a one-way ANOVA with Tukey correction (* p>0.05).

To assess potential differences in nanoparticle stability over time during storage at 4°C, physicochemical characterization was repeated for a subset of generated formulations (isomer-pure LNPs, LNPs with an equimolar cKK-E12 mixture, and commercial cKK-E12 isomer mix) over 6 weeks. Particle sizes and PDIs decreased significantly over time for all tested formulations, except for the formulation containing the commercial cKK-E12 isomer mix (Fig. 2f, g). By contrast, encapsulated mRNA content remained stable over time for *cis*-cKK-E12-LNPs but decreased for *trans*-cKK-E12-containing formulations, with the greatest effect observed for the formulation containing the commercial isomer mix (Fig. 2h). Altogether, these results demonstrate that *cis*-cKK-E12 generates more favorable LNP properties than *trans*-cKK-E12, with a clear content-dependent effect in formulations containing a mixture of the two isomer variants.

### *cis-*LNPs enable higher *in vitro* and *in vivo* protein expression than *trans-*LNPs

Given differences in physicochemical properties, we sought to assess EGFP expression levels in formulations containing isomer mixtures. Therefore, *in vitro*, DC2.4 murine dendritic cells were transfected with these formulations, and 24 h later, EGFP expression levels were visualized by confocal imaging (Fig. 3a). A clear content-dependent effect was observed, with protein expression increasing as the *cis* content increased (Fig. 3b), confirmed by flow cytometry (∼4-fold increase; Fig. S15). Subsequently, it was investigated whether this effect could be observed *in vivo*. Therefore, the same formulations were evaluated in wild-type zebrafish embryos, a convenient, cost-effective, medium- to high-throughput *in vivo* screening model that allows facile imaging-based nanoparticle biodistribution and protein expression analysis.^18–23^ EGFP expression patterns in zebrafish embryos matched the *in vitro* observations, with clear increases in protein expression as the *cis*-cKK-E12 content in the formulation increased (Figs. 3c, S16). Notably, quantification of protein expression levels in zebrafish embryos revealed that *cis-*LNPs resulted in statistically significantly higher protein expression levels than those from *trans-*LNPs and from formulations containing the commercial isomer mix (Fig. 3d).

**Fig. 3.**
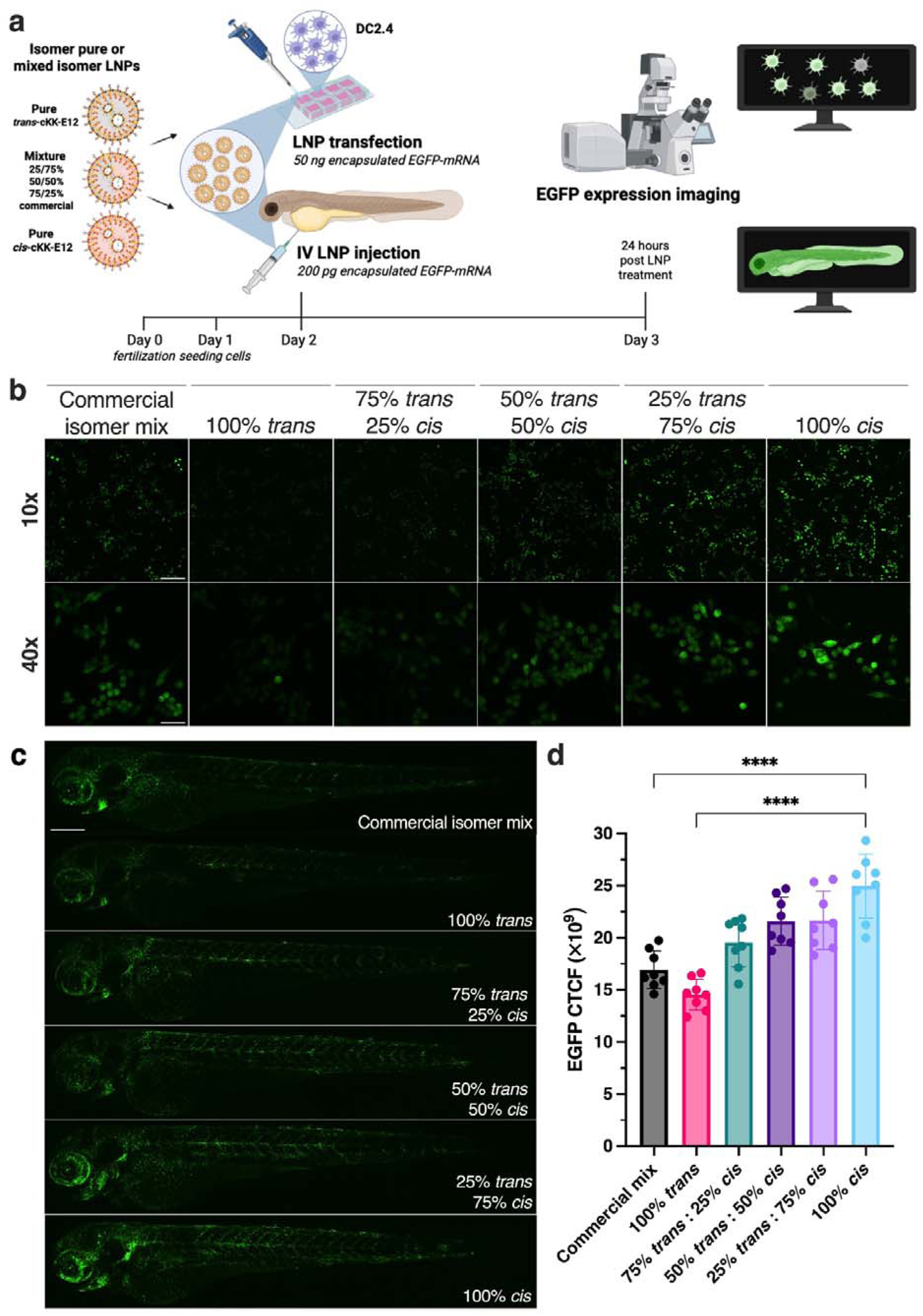
*In vitro* and *in vivo* EGFP expression analysis of cKK-E12 isomer-pure or mixed isomer lipid nanoparticle formulations. **a**, Schematic representation of the experimental procedure. DC2.4 cells were seeded in a confocal 8-well plate and treated with 50 ng EGFP-mRNA encapsulated in LNPs containing pure cKK-E12 isomers or defined isomer mixtures, as well as a formulation containing the commercial isomer mix as a reference. Wild-type AB/TL zebrafish received the same formulations at an mRNA dose of 200 pg (0.2 mg/kg). 24 h later, EGFP expression was visualized by confocal imaging. **b**, Confocal maximum intensity z-projections of DC2.4 cells 24 h after treatment with isomer pure or mixed isomer formulations captured at 10× (scale bar represents 200 µm) and zoomed in at 40× (scale bar represents 50 µm). **c**, Confocal maximum intensity z-projections of wild-type (AB/TL) zebrafish embryos 24 h after receiving 0.2 mg/kg encapsulated EGFP-mRNA in either of the formulations. Scale bar represents 250 µm. **d**, Quantification of EGFP expression levels in wild-type zebrafish embryos by corrected total cell fluorescence (CTCF). Data is presented as group mean (n=8) ± SD and statistically compared by a one-way ANOVA with Tukey correction (**** p>0.0001).

### Helper lipid isomers affect cellular uptake and protein expression of cKK-E12 LNPs *in vitro*

Driven by the clear impact of ionizable lipid isomer identity on physicochemical properties and protein expression *in vitro* and *in vivo*, it was hypothesized that helper lipid isomers would similarly influence cellular uptake and protein expression. To assess the role of lipid isomer identity in LNP behavior more comprehensively, helper lipid isomers were incorporated into isomer-pure *cis*- or *trans*-LNPs. Specifically, the DSPC enantiomer, **2,3**-distearoyl-*sn*-glycero-1-phosphocholine (Ent-DSPC), and the epimeric diastereomer of cholesterol, cholest-5-en-**3**α-ol (epicholesterol), were used to replace DSPC and cholesterol in the standard formulation. Epicholesterol, rather than Ent-cholesterol, was selected because it represents only a minor structural modification (the inversion of a single stereocenter). Additionally, DOPE was included as an alternative helper lipid to replace DSPC, as it is commonly paired with cKK-E12 and has been shown to enhance protein expression through inverse hexagonal phase formation, increasing fusogenicity.^12–16^

The effects of these compounds were screened in ten LNP formulations combining different cKK-E12 and helper lipid isomers/variants at a fixed molar ratio of 50:10:37.3:2.5:0.2 (cKK-E12:phospholipid:sterol:DMG-PEG2K:DiD) (Fig. 4a,b). Physicochemical characterization confirmed the previously observed trend that *cis*-cKK-E12 enables more efficient mRNA encapsulation at smaller particle sizes and PDIs than *trans*-cKK-E12. Helper lipid isomer identity, however, had no clear effect on particle size or PDI. Encapsulation efficiency was slightly reduced in *trans* formulations containing epicholesterol and/or Ent-DSPC compared to cholesterol and DSPC. Notably, pairing *trans*-cKK-E12 with DOPE improved mRNA encapsulation efficiency relative to DSPC or Ent-DSPC at slightly smaller particle sizes and PDIs (Fig. 4c).

**Fig. 4.**
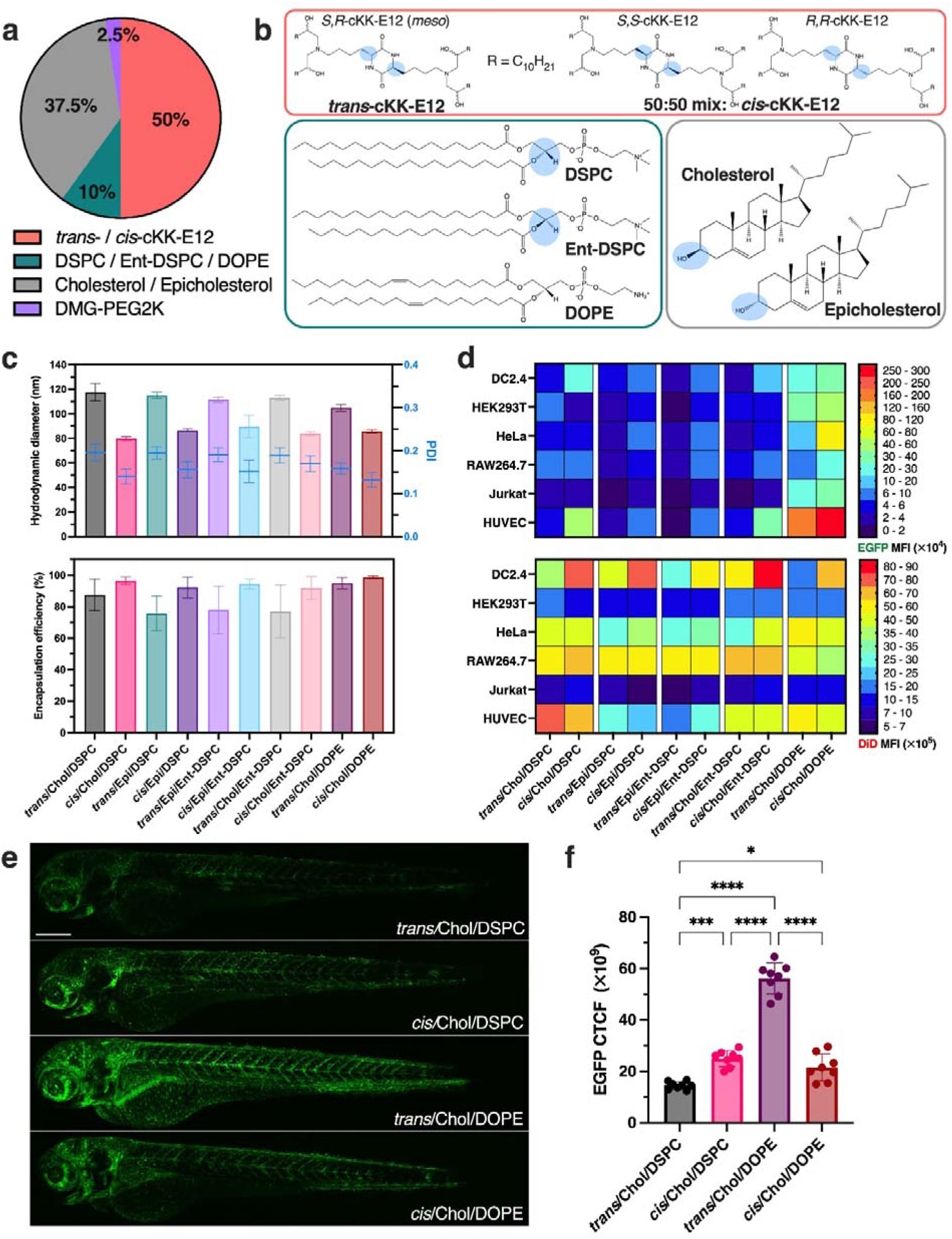
The impact of cKK-E12 and helper lipid (isomer) identity on LNP physicochemical properties and EGFP expression levels *in vitro* and in zebrafish embryos. a, Formulation design and lipid molar ratio used in these experiments. **b**, Chemical structures of lipid (isomers) used in these experiments. *R,S*-cKK-E12 (*meso*) was named *trans*-cKK-E12 and the 50:50 mixture of *R,R*- and *S,S*-cKK-E12 was designated *cis*-cKK-E12 for convenience. **c**, Particle size, PDI, and encapsulation efficiency of ten formulations containing distinct combinations of cKK-E12 and helper lipid isomers. **d**, Heatmap showing EGFP expression levels and DiD fluorescence intensities of six different cell lines (DC2.4, HEK293T, HeLa, RAW264.7, Jurkat, and HUVEC) 24 h after treatment with 500 ng/mL of EGFP-mRNA encapsulated in the ten generated formulations labeled with 0.2 mol% DiD. **e**, Confocal z-projections of wild-type (AB/TL) zebrafish embryos 24 h after receiving 0.2 mg/kg EGFP-mRNA encapsulated in DSPC- and DOPE formulations containing either *trans*- or *cis*-cKK-E12. Scale bar represents 250 µm. **f**, Quantification of EGFP expression levels in wild-type zebrafish embryos by corrected total cell fluorescence (CTCF). Data is presented as group mean (n=8) ± SD and statistically compared by a one-way ANOVA with Tukey correction (* p>0.05, *** p>0.001, **** p>0.0001).

These ten formulations were transfected into six cell lines to evaluate differences in cellular uptake and protein expression. Flow cytometry of EGFP expression and DiD fluorescence revealed clear variation in both. Uptake generally reflected endocytic capacity: phagocytes (DC2.4, RAW264.7), cancer cells (HeLa), and endothelial cells (HUVECs) showed higher uptake than low-endocytic cells (Jurkat, HEK293T). However, uptake levels varied across cell lines, with DC2.4 showing a clear preference for *cis*-LNPs compared to trans counterparts, regardless of helper lipid (isomer) identity. By contrast, other lines showed weaker (HEK293T, HeLa, RAW264.7) or helper lipid–dependent preferences (Jurkat, HUVEC). In some cases, *trans* formulations were internalized more efficiently, for example, in HEK293T and HUVECs (Chol/DSPC).

Regarding helper lipid isomer identity, formulations containing epicholesterol and/or Ent-DSPC generally showed reduced uptake compared to Chol/DSPC, particularly when both were combined, although the extent varied across cell lines. This reduction was most pronounced in HUVECs and, to a lesser extent, Jurkat cells, which exhibited lower uptake of Epi/DSPC and Epi/Ent-DSPC. Additionally, DSPC- and Ent-DSPC-containing formulations were better internalized than DOPE formulations in some cell lines, especially in DC2.4 and, to a lesser extent, HUVECs (Chol/DSPC). Notably, across all cell lines, *trans*/Epi/Ent-DSPC consistently ranked among the least efficiently internalized formulations (Figs. 4d, S17).

Regarding EGFP expression, higher levels were generally observed for DOPE formulations compared to all others, consistent with previous reports^13,15,16^. The most pronounced effect was observed in HUVECs, which internalized *trans*/Chol/DSPC most efficiently but exhibited 51-fold higher EGFP expression for *cis*/Chol/DOPE. Overall, EGFP expression broadly correlated with cellular uptake, with notable exceptions in HUVECs for *trans*/Chol/DSPC and *cis*/Epi/DSPC. Although the magnitude of the differences was cell-line-dependent, *cis* formulations generally outperformed their *trans* counterparts. An exception was *trans*/Chol/DSPC in HEK293T, which exceeded *cis*/Chol/DSPC, likely due to differences in uptake. Differences between *cis* and *trans* formulations were most pronounced in DC2.4 and HUVECs and smallest in HeLa, Jurkat, and RAW264.7. In these latter cell lines, *cis*–*trans* differences were greater for formulations containing epicholesterol and/or Ent-DSPC than for Chol/DSPC. Overall, *trans*/Epi/Ent-DSPC performed worst, while *cis*/Chol/DOPE performed best across all tested cell lines (Figs. 4d, S17). Cytotoxicity was evaluated in HEK293T cells and was only substantially increased for the cis/Chol/DOPE formulation (Fig. S17c).

Altogether, *cis-*LNPs performed better than their *trans* counterparts, regardless of the helper lipid (isomer) identity. Additionally, Chol/DSPC formulations were generally internalized more efficiently and were associated with better EGFP expression than Epi/Ent-DSPC formulations. These results indicate a profound effect of both the stereochemistry of ionizable and helper lipids on lipid nanoparticle behavior *in vitro*.

Notably, the two DOPE formulations were also injected into wild-type zebrafish embryos to evaluate potential differences in protein expression *in vivo*. Interestingly, the reverse trend was observed compared to the *in vitro* results, with *trans*/Chol/DOPE outperforming *cis*/Chol/DOPE (Fig. 4e,f).

### *cis*-LNPs’ protein expression levels significantly outperform *trans*- equivalents in mice

The clear *in vitro* finding that *cis*-LNPs generally outperform *trans* counterparts prompted evaluation in a rodent model. The ten previously developed formulations, combining different cKK-E12 and helper lipid isomers, were therefore reformulated to encapsulate firefly luciferase (FLuc) mRNA for *in vivo* bioluminescence imaging. In addition, a reference formulation containing the commercial cKK-E12 isomer mix paired with DSPC and cholesterol was included. Mice received 0.15 mg/kg FLuc-mRNA encapsulated in either of these eleven formulations, followed by intravenous D-luciferin administration 6 h later (Fig. 5a).

**Fig. 5.**
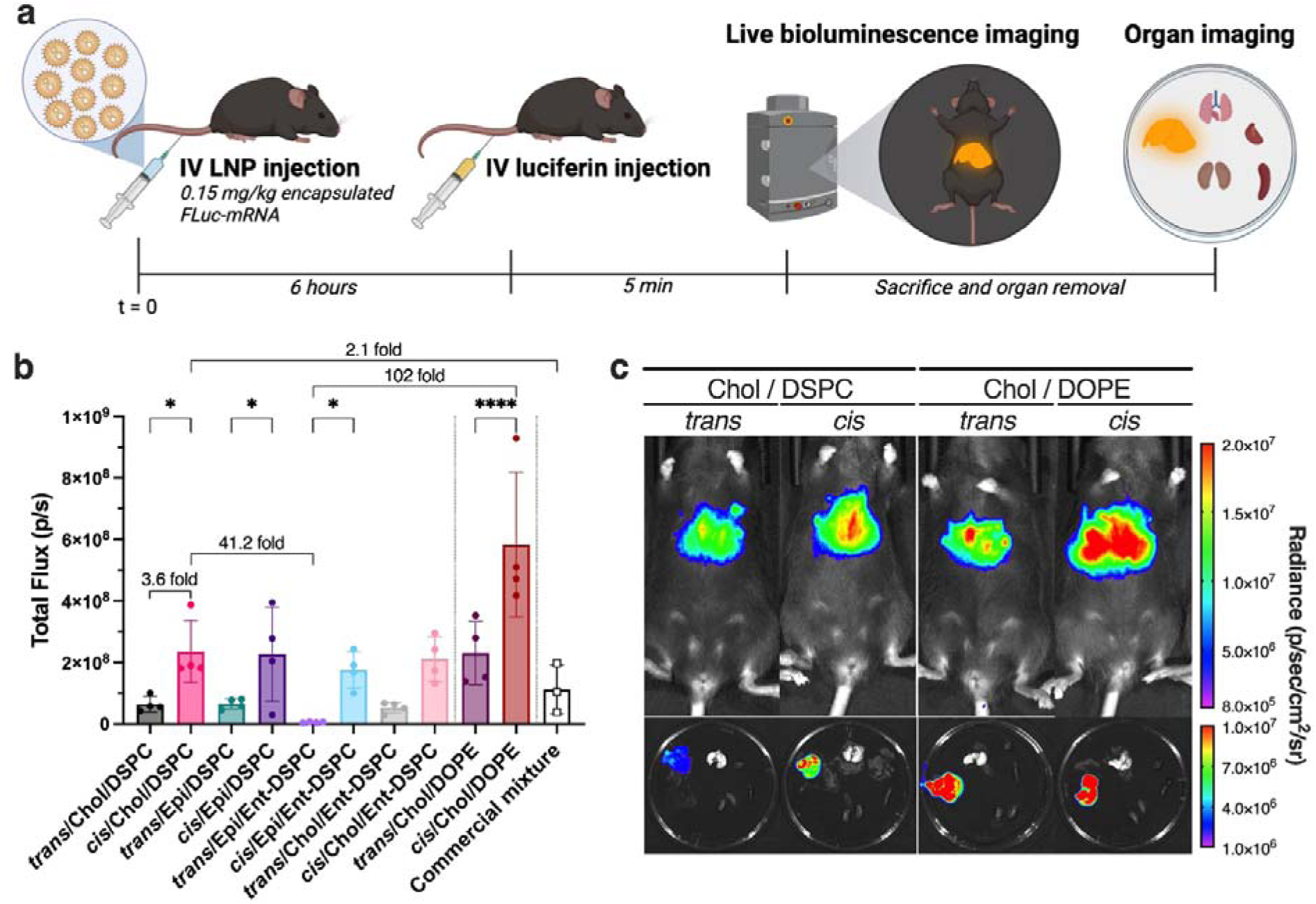
*cis* formulations significantly outperform *trans* equivalents regardless of helper lipid (isomer) identity in mice. **a**, Schematic representation of the experimental procedure. C57BL/6 mice received (IV) 0.15 mg/kg FLuc-mRNA encapsulated in one of the ten developed formulations, each combining cKK-E12 with different helper lipid isomers, or the reference formulation containing the commercial cKK-E12 isomer mix. 6 h later, D-luciferin was administered (IV), followed by live bioluminescence imaging using IVIS after 5 min. Subsequently, mice were sacrificed, and the liver, spleen, lungs, kidneys, and heart were removed from one animal per group and imaged using IVIS to determine biodistribution. **b**, Quantification of FLuc expression by total flux per second. Data are presented as group mean (n=4) ± SD and statistically compared by one-way ANOVA with Tukey correction (* p>0.05, **** p>0.0001). c, Live bioluminescence IVIS images of representative individuals that received *trans*/Chol/DSPC, *cis*/Chol/DSPC, *trans*/Chol/DOPE, or *cis*/Chol/DOPE formulations. Organ images do not necessarily originate from the same animals as the live images and were used only for general biodistribution analysis.

IVIS imaging revealed clear differences between *cis*- and *trans* formulations, independent of helper lipid isomer identity. DSPC-containing formulations slightly outperformed those with Ent-DSPC, although differences were not significant. Notably, *trans*/Epi/Ent-DSPC performed markedly worse than all other trans formulations. Consistent with *in vitro* data and previous reports^13,15,16^, DOPE formulations significantly outperformed those containing DSPC or Ent-DSPC. For example, *trans*/Chol/DOPE performed similarly to *cis* formulations with DSPC or Ent-DSPC, while *cis*/Chol/DOPE clearly outperformed all others. The reference formulation containing the commercial isomer mix performed slightly better than pure *trans* formulations, but remained below *cis* formulations, although not significantly (Fig. 5b). Interestingly, only mice receiving the reference formulation showed bioluminescence on palmar and, to a lesser extent, plantar pads, a pattern not observed with any other formulation (Fig. S18a).

Mice were subsequently sacrificed, and the liver, spleen, lungs, kidneys, and heart from one animal per group were collected for IVIS-based biodistribution analysis. Consistent with previous reports^12,24^, all formulations primarily targeted the liver (Fig. S18b), with expression levels reflecting trends observed in live imaging. For one animal treated with *cis*/Chol/DOPE, the liver was removed, and the remaining organs reimaged, revealing detectable FLuc expression only in the spleen (Fig. S18c). Quantification showed liver expression was 62-fold higher than in the spleen (Fig. S18d).

Overall, these *in vivo* observations are consistent with some of the *in vitro* results, particularly those obtained with HUVECs and DC2.4, and underscore the significant impact of ionizable and helper lipid stereochemistry on *in vivo* LNP performance.

### Internal lipid organization underlies *cis*–*trans*-cKK-E12 efficacy differences

With these clear differences in efficacy, both *in vitro* and *in vivo,* in mind, we sought to investigate a possible structure-function relationship that could explain these observations. Therefore, four formulations (*trans*/Chol/DSPC, *cis*/Chol/DSPC, *trans*/Chol/DOPE, and *cis*/Chol/DOPE) were selected for in-depth structural characterization using AI-assisted cryogenic transmission electron microscopy (cryo-TEM) and small-angle X-ray scattering (SAXS) (Fig. 6).

**Fig. 6.**
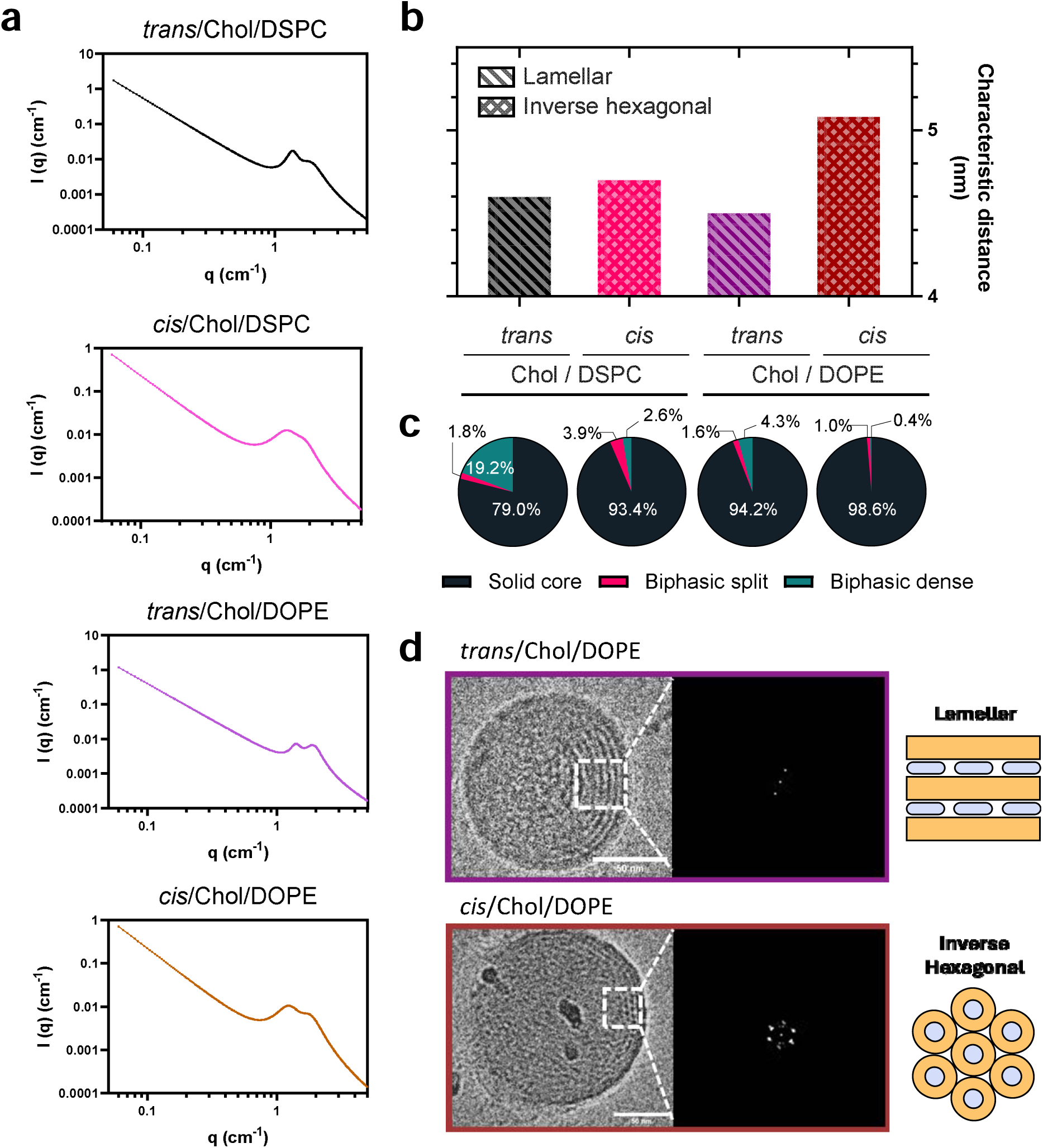
cKK-E12 and helper lipid isomer identity influence internal lipid organization of LNPs. **a**, SAXS profiles of *trans*/Chol/DSPC, *cis*/Chol/DSPC, *trans*/Chol/DOPE, and *cis*/Chol/DOPE formulations loaded with EGFP-mRNA. **b,** Characteristic distance of *trans*/Chol/DSPC, *cis*/Chol/DSPC, *trans*/Chol/DOPE, and *cis*/Chol/DOPE formulations, as determined from the SAXS profiles (first peak) with classification as hexagonal or lamellar as determined by Cryo-TEM. **c**, Particle morphology distributions of *trans*/Chol/DSPC, *cis*/Chol/DSPC, *trans*/Chol/DOPE, and *cis*/Chol/DOPE formulations identified by AI-assisted cryogenic transmission electron microscopy (cryo-TEM) data analysis. **d**, Magnified cryo-TEM images of *trans*/Chol/DOPE and *cis*/Chol/DOPE formulations, with corresponding fast Fourier transforms (FFTs) of selected regions and schematic illustrations representing the observed internal structures.

Regarding internal lipid organization, regions of interest in cryo-TEM images were subjected to fast Fourier transform (FFT), revealing that most *cis* formulations have inverse hexagonal (HII) internal structures, whereas *trans* formulations exhibit lamellar (Lα) organization. Notably, the *cis*/Epi/DSPC formulation displayed cubic-phase organization (Figs. 6d, S24, Table 1).

SAXS curves showed two different peaks for the studied formulations, suggesting coexistence of two distinct structural features (Figs. 6a, S19). The absence of second and third order Bragg reflections hindered the accurate structure assignment based on such peaks, and such absence is likely due to a relatively low degree of crystallinity or a rather short range of order persistence (domain size) in space. Previous studies suggest that SAXS peaks in this region can correspond to ionizable lipid-mRNA packed assemblies, including lamellar, hexagonal, and worm-like inverse micellar systems.^25–27^ The first peak, representing a characteristic distance (*d)* around 4.4-4.8 nm, generally displayed higher amplitude (*A)* than the second peak, which presented *d* values around 3.0-3.4 nm, suggesting that the former is the most dominating of the two features (Table S1). Previously, Pattipeiluhu *et al* reported larger characteristic distances when LNPs are loaded with RNA as opposed to containing only lipids.^34^ Therefore, one hypothesis for the presence of two first order Bragg peaks is the coexistence of internal structures composed of both lipids and RNA (first peak) and only lipids (second peak) in the same sample. Changing from *trans* to *cis* formulations produced a small increase in the *d* values of the first peak and a small decrease in the correlation length (ξ) of the first peak (Fig. S22). It also led to a small decrease in the amplitude of the second peak. Switching from cholesterol to epicholesterol produced a decrease in *A* and ξ values of the first peak and also a decrease in the *d* values of the second peak. Regarding the helper lipid, replacing DSPC by either Ent-DSPC or DOPE led mainly to a decrease in the *A* values of the first peak. Notably, the worst-performing formulation (*trans*/Epi/Ent-DSPC) showed the smallest *d*-value for the first peak, whereas the best-performing formulation (*cis*/Chol/DOPE) had the largest one (Table S1).

AI-assisted cryo-TEM classified LNP morphology into three types: solid core, biphasic split, and biphasic dense (blebbing). All formulations predominantly formed spherical solid-core particles, but *trans*/Chol/DSPC showed a notable ∼20% fraction of biphasic dense particles, compared to <5% for the other formulations. *trans*/Chol/DOPE displayed a more uniform morphology (>94% solid core), with the remaining fraction mainly biphasic dense (4.3%). In contrast, *cis*/Chol/DSPC, with a similar solid-core fraction, showed slightly more biphasic split particles. Notably, *cis*/Chol/DOPE consisted almost entirely of solid-core LNPs, with only 0.4% biphasic dense particles (Figs. 6c, S23).

Furthermore, apparent pK_a_ of *cis*- and *trans*-LNPs containing DSPC and Chol were determined by 2-(p-toluidino)-6-naphthalene sulfonic acid (TNS) assay and did not substantially differ at 6.6 and 6.4, respectively (Fig. S20).

Thus, these combined analyses identified internal lipid organization as the underlying cause of efficacy differences between *cis*- and *trans* formulations. In addition to lipid arrangements, characteristic distance was found to be an important factor determining LNP performance.

## Discussion

Here, we comprehensively examined how lipid stereochemical identity influences the physicochemical properties of LNPs and the resulting protein expression levels in both *in vitro* and *in vivo* settings. Initially, we observed significant effects on physicochemical properties (Fig. 2), followed by differences in protein expression among formulations with different ratios of ionizable lipid isomers (Figs. 3, S14-16), showing that the *cis* isomer outperforms the *trans* isomer. Subsequently, LNP uptake *in vitro* and protein expression differences *in vitro* and *in vivo* were greatly influenced by isomers of ionizable lipid and helper lipids (Figs. 4,5).

Interestingly, *trans*-LNPs showed higher protein expression than *cis*-LNPs containing DOPE in zebrafish embryos across two independent experiments (Fig. 4), deviating from trends observed *in vitro* (Figs. 4, S17) and in mice (Fig. 5). This may be due to differences in protein corona composition and/or abundance, which can affect serum stability, cellular uptake, trafficking, and cytoplasmic delivery^28–32^. Although adult zebrafish share similar serum protein profiles, particularly in abundant proteins such as apolipoproteins and complement proteins^33^, the serum composition of embryos remains unknown due to limited sample availability and may differ substantially given rapid developmental changes. Alternatively, higher cellular uptake of *trans* formulations, as observed *in vitro* across several cell lines (HeLa, RAW264.7, HUVEC, and to a lesser extent HEK293T), may also contribute (Figs. 4, S17).

Furthermore, besides protein expression analyses, in-depth morphological and structural analyses by cryo-TEM and SAXS revealed impacts of both ionizable and helper lipid stereochemistry on overall LNP morphology, internal lipid organization (L_α_ versus H_II_), as well as characteristic distance (Figs. 6, S24). SAXS data indicated that changes in the isomeric identity of the LNPs, including ionizable lipid, cholesterol and helper lipid can all lead to changes in the internal organization of the particles. Importantly, apparent pK_a_ of *cis*- and *trans*-LNPs (containing Chol and DSPC) did not substantially differ (Fig. S20), suggesting differences in internal lipid organization as the leading cause of mRNA delivery potency differences.

Altogether, these results demonstrate the importance of the stereochemical identity of LNPs by establishing, for the first time, structure-function relationships among lipid stereochemistry, internal lipid organization, LNP morphology, and protein expression. Overall, *cis*-cKK-E12 paired with DOPE generated H_II_ phases with the largest observed characteristic distance among the tested formulations (Fig. 6), leading to significantly higher *in vivo* efficacy than its *trans* counterpart. Notably, the worst-performing formulation (*trans*/Epi/Ent-DSPC), which had the shortest observed characteristic distance, showed over 100-fold lower protein expression *in vivo* compared with the others (Figs. 5, S22). These findings further contribute to structure-driven rational design of highly efficacious LNP formulations for mRNA delivery.

## Materials & Methods

### Reagents

cKK-E12 was purchased from Echelon Biosciences (Salt Lake City, UT, USA). 1,2-distearoyl-sn-glycerol-3-phosphocholine (DSPC), 2,3-distearoyl-sn-glycero-1-phosphocholine (Ent-DSPC), 1,2-dioleoyl-sn-glycero-3-phosphoethanolamine (DOPE), and 1,2-dimyristoyl-rac-glycero-3-methoxypolyethylene glycol-2000 (DMG-PEG2K) were purchased from Avanti Polar Lipids (Alabaster, USA). Cholesterol (14606-100G-F) was purchased from Merck KGaA (Darmstadt, Germany). Epicholesterol was obtained from Aaron Chemicals (San Diego, CA, USA). The lipid-dye conjugate DiD (1,1’-Dioctadecyl-3,3,3’,3’-Tetramethylindodicarbocyanine, 4-Chlorobenzenesulfonate Salt) was purchased from Fisher Emergo B.V. (Landsmeer, The Netherlands). TriLink CleanCap® 5-methoxyuridine (5moU) modified EGFP-mRNA and CleanCap® AG Reagent were purchased from Tebubio (Heerhugowaard, The Netherlands). Quant-IT™ RiboGreen RNA Assay Kit, 0.5 M Citrate Buffer (pH 4.0), Invitrogen reagents for IVT reactions, IVT purification, and agarose gel staining, and Gibco supplements for culture media were purchased from Fisher Emergo B.V. (Landsmeer, The Netherlands). DPBS (10×) was obtained from VWR International B.V. (Amsterdam, The Netherlands). Vivaspin 500 centrifugal filters (MWCO 100K or 300K) were obtained from Sartorius (Göttingen, Germany). All primers for PCR were obtained from Integrated DNA Technologies B.V. (Leuven, Belgium). New England Biolabs (NEB) reagents for *in vitro* transcription (IVT) and DNA or RNA purification were purchased from Cell Signaling Technology Europe B.V. (Leiden, The Netherlands). RPMI and DMEM culture media and fetal bovine serum were obtained from Capricorn Scientific GmbH (Ebsdorfgrund, Germany). PromoCell Endothelial Cell Growth Medium MV2, WST-1 cell viability reagent, and ampicillin (sodium salt; A8351) were purchased from Merck KGaA (Darmstadt, Germany). Thermo Scientific™ Pierce™ D-Luciferin, Monopotassium Salt was purchased from Life Technologies Corporation (Bleiswijk, The Netherlands).

### cKK-E12 isomer characterization and purification

cKK-E12 was characterized by analytical HPLC equipped with an ELSD detector using a C18 column (Phenomenex Gemini^®^, 3 μm, C18, 110 Å, 50 x 4.6 mm). The *cis-trans* mixture of cKK-E12 was purified by preparative HPLC with column oven and ELSD detector, employing a C18 column (Phenomenex Kinetex^®^, 5 μm, EVO C18, 100 Å, 150 x 21.2 mm) and a of 30-90% acetonitrile/water gradient containing 0.1% TFA ACN/H_2_O with 0.1% TFA at 50 °C. After isolation the *trans* and *cis* fractions were freeze dried to yield *cis*-cKK-E12 and *trans*-cKK-E12 as a white powder.

### Lipid nanoparticle manufacturing

Lipid nanoparticles were self-assembled as previously described^34^ with minor adaptations. Lipids were combined at the desired molar ratios and concentrations from stock solutions (0.05–10=mM) in chloroform. The molar ratio between the lipids used in this study was 50:10:37.3:2.5:0.2 (cKK-E12:phospholipid:cholesterol:DMG-PEG2K:DiD). Mixtures of purified cKK-E12 isomers always constituted 50 mol% of the total lipid mixture. Solvents were evaporated under a nitrogen flow. The lipid film was dissolved in absolute ethanol to a total lipid concentration of 4–6 mM. 1 mg/ml RNA stored at -80°C was thawed and diluted in 25=mM citrate buffer to a final mRNA concentration of 33.33–50 µg/mL. Depending on the amount of mRNA to be encapsulated, volumes ranged from 150–300 µL and 450–900 µL for the lipid solution and the mRNA solution, respectively. The solutions were loaded into two separate sterile syringes, connected to a T-junction microfluidic mixer, and mixed using two Chemyx Fusion 100X syringe pumps at a 3:1 (RNA:lipid) flow ratio, yielding a total flow rate of 2.4 mL/min. After mixing, the solution was directly loaded into a dialysis cassette (Slide-A-Lyzer™ 20 kDa MWCO 0.5 mL, Thermo Scientific) and dialyzed against PBS for 2–16 hours. LNPs were concentrated using Vivaspin 500 centrifugal filters at 2000 × g, 4 °C.

### Lipid nanoparticle characterization

LNP sizes and zeta potentials were measured using a Malvern Zetasizer Nano ZS (software version 7.13, Malvern Panalytical). For DLS (operating wavelength = 633 nm), measurements were performed at 25 °C in 1x PBS (pH = 7.4) at a total lipid concentration of approximately 0.1 mM. Zeta potentials were measured at 0.5 mM total lipid concentration, using a folded capillary cell (DTS1070, Malvern), at room temperature. All reported DLS measurements and zeta potentials are the average of three measurements.

Encapsulation efficiency and encapsulated RNA concentration were analyzed using the Quant-iT^TM^ Ribogreen RNA assay by diluting 1–2 µL of concentrated LNPs back to their original concentration after dialysis in 1x PBS, and then following the manufacturer’s low-range assay protocol. RNA encapsulation efficiency was calculated using the following equation:

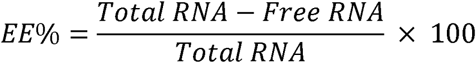

### CD measurements

CD spectra were recorded on a JASCO J-815 CD spectrometer equipped with a Peltier temperature controller. Samples were measured at 20 °C in a quartz cuvette with a 2 mm path length. Spectra were aquired from 200 to 300 nm at 0.5 nm intervals using a 1 nm bandwidth. Each final spectrum represented the average of five sequentially recorded scans. Samples were measured at a concentration of 400 µM in acetonitrile. The mean residue molar ellipticity (θ, deg·cm2·dmol-1) was calculated according to the following equation:

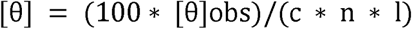

### Firefly luciferase mRNA design, *in vitro* transcription, and purification

A gBlock DNA fragment containing a *Mus musculus* iCodon-optimized^35^, red-shifted firefly luciferase coding sequence (amino acid sequence derived from a previous report^36^) was obtained from Integrated DNA Technologies (Coralville, IA, USA). A pUC57-mini vector containing a T7 promoter with AGG initiator, synthetic 5’ untranslated region (UTR)^37^, human beta globin 3’ UTR^38^, and poly(A_35_) sequence inserted at the EcoRV site was obtained from GenScript Biotech B.V. (Rijswijk, The Netherlands).

The pUC57-mini vector was linearized by PCR with primers that introduce 20 nt overlaps with the FLuc DNA fragment, using Q5^®^ High-Fidelity Hot Start DNA polymerase (New England Biolabs, NEB), and purified with the Monarch^®^ PCR and DNA Cleanup kit (NEB). Subsequently, the FLuc DNA fragment was cloned into the linearized vector using NEBuilder^®^ HiFi DNA assembly master mix (NEB) and transformed into Stable Competent *E. coli* (NEB). A PCR-prescreened clone was grown overnight in LB supplemented with ampicillin (100 µg/mL) to amplify the desired plasmid. The plasmid was isolated using the Monarch^®^ Plasmid Miniprep kit (NEB). Subsequently, the sequence of the complete FLuc construct (included in Fig. S17) was verified by Sanger sequencing performed by BaseClear (Leiden, The Netherlands).

Template for *in vitro* transcription (IVT) was generated by PCR using Q5^®^ High-Fidelity Hot Start DNA polymerase (NEB) in a 100 µL reaction using a reverse primer annealing at the 3’ end of the 3’ UTR, installing a poly(A_120_) tail. Then, the size and purity of the PCR-produced IVT template were verified by a 1.2% agarose gel stained with SYBR^TM^ Safe (Invitrogen). Subsequently, any circular plasmid DNA used as the template in the PCR reaction was digested by adding 2 µL of DpnI (NEB) and incubating for 20 minutes at 37°C in a thermal cycler. Finally, the IVT template was purified using the Monarch^®^ PCR and DNA Cleanup kit (NEB), and the yield was measured using NanoDrop One (Thermo Fisher Scientific).

FLuc-mRNA was synthesized by *in vitro* transcription using the following reaction conditions: 40 ng/µL purified IVT template, 3.75 U/µL Hi-T7 RNA polymerase (high concentration; NEB) accompanied with 1× of its supplied buffer, 5 mM ribonucleotides in which UTP was replaced with 5moUTP (NEB), 1 U/µL Murine RNase inhibitor (NEB), 4 U/mL Yeast inorganic pyrophosphatase (NEB), 4 mM CleanCap^®^ AG Reagent (TriLink BioTechnologies), 26 mM MgCl_2_ solution, and 5 mM DTT solution (both Invitrogen). The reaction was incubated for 3 hours at 50°C in a thermal cycler. Afterwards, 1 µL of DNase I-XT (NEB) per µg of template DNA was added to the reaction and incubated for 15 min at 37 °C in a thermal cycler. Immediately thereafter, the IVT reaction was purified using the Monarch^®^ Spin RNA Cleanup kit (500 µg; NEB) and eluted into 1 mM citrate buffer, pH 6.4 (Invitrogen). The integrity and purity of 100 ng of purified IVT mRNA were verified using a 2% agarose gel stained with SYBR^TM^ Gold (Invitrogen), along with the High Range ssRNA ladder (NEB), both diluted in the supplied 2× RNA loading dye, following the manufacturer’s sample preparation recommendations. The IVT mRNA was diluted to 1 mg/ml concentration, snap-frozen in dry ice and stored at -80 °C until use.

### Cell culture and mRNA-LNP transfection

The murine immortalized dendritic cell line DC2.4 was kindly provided by Kenneth Rock, University of Massachusetts Medical School, Worcester, MA, USA. The murine immortalized macrophage cell line RAW264.7 and the human cell lines HEK293T, HeLa, and Jurkat were obtained from ATCC^®^ (VA, USA). HUVEC cells (C-12203) were purchased from Merck KGaA (Darmstadt, Germany). DC2.4 and Jurkat cells were cultured in RPMI 1640 without L-Glutamine. HEK293T, HeLa, and RAW264.7 were cultured in DMEM High Glucose (4.5 g/L) without L-Glutamine. Both media were supplemented with 10% Advanced heat-inactivated FBS, 1% penicillin-streptomycin, and 2 mM GlutaMAX™. HUVEC cells were cultured in Endothelial Cell Growth Medium MV 2 (PromoCell) supplemented with the supplied supplement mix. Cells were maintained at 37 °C in a humidified atmosphere with 5% CO_2_.

### *In vitro* mRNA-LNP uptake and expression analysis

The mRNA-LNP transfection and expression studies were performed using EGFP-mRNA-LNPs labeled with 0.2 mol% DiD. Briefly, cells—DC2.4, RAW264.7, HEK293T, HeLa, Jurkat, and HUVEC—were seeded at 1–2 × 10^4^ cells/well in a 96-well flat-bottom plate and allowed to adhere for 24 hours. Cells were then treated with encapsulated EGFP-mRNA at concentrations of 100, 250, and 500 ng/mL in supplemented medium and incubated for 24 hours at 37 °C in a 5% CO_2_ atmosphere. After incubation, cells were washed with PBS, resuspended in FACS buffer (PBS, 2% FBS, 0.1% sodium azide, and 1 mM EDTA), and analyzed by flow cytometry (CytoFLEX S, Beckman Coulter, CA, USA) for DiD and EGFP fluorescence. Data were analyzed using FlowJo software V10 (Tree Star Inc., OR, USA). The mean fluorescence intensity (MFI) of EGFP and DiD was graphed using GraphPad Prism 10.

For confocal imaging, DC2.4 cells were seeded in an 8-well plate (µ-Slide, Ibidi) at 4 × 10^4^ cells/well and incubated at 37 °C for 24 hours. Subsequently, cells were treated with PBS or DiD-labeled EGFP-mRNA-LNPs at an mRNA concentration of 500 ng/mL and incubated for 24 hours. After treatment, cells were washed with PBS, stained with Hoechst 33342 for nuclear visualization, incubated for 30 minutes at 37 °C, and imaged immediately thereafter using a Leica TCS SP8 confocal laser scanning microscope (Leica Microsystems, Wetzlar, Germany). Confocal Z-stacks were acquired using a Leica TCS SP8 confocal laser-scanning microscope, with either a 10× air objective (HCX PL FLUOTAR) or a 40× water-immersion objective (HCX APO L) for zoom-ins. To ensure comparability across experiments, microscopy settings, such as laser intensity, gain, and offset, were kept constant between stacks. Maximum-intensity Z-projections of individual and merged channels were generated using FIJI imaging software^39^.

### Cell viability assay

HEK293T cells were seeded in 96-well plates at a density of 2 × 10^4^ viable cells/well and allowed to attach for 24 hours at 37 °C. The cells were treated with EGFP-mRNA-LNP formulations at different mRNA concentrations (10–2500 ng/mL). 24 hours later, cells were washed with PBS, and cell viability was determined by the WST-1 cell viability assay. Viability of LNP-treated cells was normalized to that of untreated cells, with 100% viability.

### Animal experiments and husbandry

All animal work was approved by the Leiden University Animal Ethics Committee and the animal experiments were performed conform the guidelines from Directive 2010/63/EU of the European Parliament on the protection of animals used for scientific purposes. Wild-type C57Bl/6 mice were purchased from Jackson Laboratory (CA, USA), bred in-house, kept under standard laboratory conditions, and provided with food (chow) and water *ad libitum*.

Zebrafish were maintained in accordance with standard zebrafish rearing protocols (https://zfin.org), which adhere to the international guidelines of the EU Animal Protection Directive 2010/63/EU. Housing and husbandry recommendations were followed as previously recommended^40^. Fertilization was performed by natural spawning at the beginning of the light period, and eggs were raised at 28°C in egg water (60 μg/ml Instant Ocean sea salts). Only wild-type AB/TL zebrafish were used in this study.

### Zebrafish microinjections confocal imaging, and image processing

LNP formulations were injected into zebrafish embryos at time points ranging from 54–60 hours post-fertilization (hpf) using a modified microangraphy protocol^41^. One nanoliter volume of nanoparticle formulation was calibrated and injected into the duct of Cuvier after the embryos were embedded in 0.4% agarose containing 0.01% tricaine as described^18^.

At least three randomly selected, well-injected embryos were mounted in a 60 mm confocal dish with 0.4% agarose in egg water containing 0.01% buffered 3-aminobenzoic acid ethyl ester (tricaine; Sigma-Aldrich, A-5040) and imaged at 24 hours post-injection (hpi). Eight whole embryos were imaged per group.

Confocal Z-stacks were captured using a Leica TCS SP8 confocal laser-scanning microscope, equipped with a 10× air objective (HCX PL FLUOTAR) for whole-body imaging. To ensure comparability across experiments, microscopy settings, including laser intensity, gain, and offset, were kept constant across stacks and sessions. Whole-embryo images consisted of 3–4 overlapping Z-stacks stitched using FIJI imaging software^39^.

Whole-embryo EGFP CTCF was measured using FIJI (ImageJ) by selecting the sum of slices’ Z-projection of the entire embryo with the wand tool, then measuring MFI, area, and integrated density of the selected region, along with background fluorescence MFI outside the embryo. CTCF was calculated, and background fluorescence was subtracted by computing (integrated density) – (area × background MFI).

### *In vivo* luciferase expression and IVIS imaging

12-week-old female mice (n = 4 per group; n = 3 for the control group with commercial cKK-E12) received 0.15 mg/kg of encapsulated FLuc-mRNA intravenously via tail vein injection in a 100 µL volume. Six hours later, they received a second 100 µL tail vein injection of a 3 mg/mL D-luciferin solution in Dulbecco’s PBS (150 mg/kg) and were allowed to recover for 3 minutes. Subsequently, mice were anesthetized with isoflurane in oxygen (induction 3%, maintenance 2%) and imaged in a light-tight IVIS system on a warmed stage at 5 minutes post-D-luciferin injection, with a constant 5-second exposure time. Afterwards, animals were sacrificed by cervical dislocation, and organs (liver, spleen, lungs, kidneys, and heart) of one animal per experimental group were removed, placed on a 9 cm petri dish, and imaged using a 20-second constant exposure time. Mice that received LNPs but no D-luciferin, as well as the IVIS chamber without mice, were imaged under the same conditions. No difference in background signal was detected between the two images; therefore, no background signal was subtracted from any image.

Live bioluminescence images were analyzed using the PerkinElmer Living Image software (version 4.7.3). The radiance (photons/sec/cm^2^/sr) scale was adjusted across all images to the range 8×10^5^–2×10^7^ for comparison. ROIs were selected using a fixed threshold of 10 or manually adjusted to organ size, since the resulting ROI size was often not representative of the organ size. ROIs at the injection site were ignored when below 5×10^6^ or added to the other ROI value for that individual. Overall data trends were the same for both ROI selection methods, thus the manual ROI selection and threshold setting was employed as it better reflected organ size and differences between groups. ROI values were measured and exported, and total flux per second was plotted using GraphPad Prism 10. Organ images were analyzed similarly, with an adjusted radiance scale of 1×10^6^ – 1×10^7^ or 5×10^5^ – 1×10^7^, but not used for quantification. For one animal that received *cis*-cKK-E12/cholesterol/DOPE, the organs were imaged with and without the liver, and the radiance scale was adjusted to 5×10^4^ – 1×10^6^.

### General Cryo-electron microscopy

Freshly glow-discharged carbon grids supported on Cu (Lacey carbon film, 200 mesh, Electron Microscopy Sciences, Aurion, The Netherlands) were used for vitrification inside a Vitrobot plunge-freezer (FEI VitrobotTM Mark III, Thermo Fisher Scientific), which regulated steady temperature and humidity conditions (22°C and 99% humidity). Concentrated mRNA-LNP formulations (4 µL, 10-15 mM) were applied to a grid, blotted for 3 seconds at 99% humidity. Cryo-TEM images were collected on a Glacios 2 operating a FEG at 200 keV, with the working temperature maintained below −180 °C. Images were acquired with Smart EPU software at a nominal magnification of 79 000x (pixel size = 1.6 ångström (Å)) and a nominal underfocus of -2 µM.

### Cryo-EM at ATEM (AI-assisted)

#### Sample Preparation

Refrigerated samples were kept on ice. 4 µL of the undiluted sample was applied to ATEM regularly spaced holey carbon copper grid (ATEM LNPCFoil Grids) in H_2_O saturated atmosphere at 4 °C and incubated for 1 min in a ThermoFisher Vitrobot Mark IV System. Excess sample was blotted away, leaving a several 100 nm thin layer of liquid on the surface of the grid. Blotted grids were vitrified immediately by plunge-freezing in liquid ethane at -180 °C. Samples were stored in sample specific containers under liquid nitrogen (lN_2_) until further use in the cryo-electron microscope.

#### Cryo-Electron Microscopy

After grid preparation, all grids were processed for transfer into the microscope and evaluated in a 200 kV ThermoFisher Glacios cryo-transmission electron microscope (cryo-TEM) equipped with a Falcon IVi direct electron detector and ThermoFisher EPU software. A low magnification (150 x magnification) overview image (Atlas) of each cryo-grid was prepared. Each Atlas of every grid was evaluated by a trained operator and evaluated for ideal ice thickness, paying particular attention to identify areas with an approximate ice thickness of 200-300 nm. Suitable areas for image acquisition and appropriate defocus settings were defined manually by a trained operator within the areas of ideal ice thickness. Image acquisition at approx. 45,000 x magnification (on specimen level) was initiated. All data was recorded at low-dose conditions (total dose: 30 e- / Å2) on a Falcon IVi direct electron detector camera using dose fractionation (10 frames). Dose fractionated raw data images (micrographs) were then processed and frame-aligned to compensate for beam induced motion and restore fine details of the imaged sample. Frame-aligned and computationally summed images were stored for data further evaluation.

#### Data Evaluation & Analysis

All preprocessed micrograph data was analyzed by a versioned, ATEM proprietary Machine Learning (ML) model. The model is trained and validated to automatically and reliably detect individual LNP particles on typical cryo-EM micrographs, extract morphological parameters like particle area, Particle perimeter and inferred diameter, particle shape. All particles are furthermore classified into one of three morphology classes (Solid Core, Biphasic Dense (i.e. “Blebbed”) or Biphasic Split). Image artifacts and particles that do not express morphological properties of either class are not evaluated. Relevant numerical and image data is extracted accordingly. Samples from each analysis are reviewed by a trained operator for accuracy and consistency for final quality control.

### TNS binding assay

The apparent pK= of LNPs was determined using the TNS binding assay. A 300 μM stock solution of TNS in DMSO was prepared and diluted together with LNPs to a final concentration of 75 μM ionizable lipid and 6 μM TNS in a total volume of 93 μL of buffered solution. The buffer contained 20 mM boric acid, 10 mM imidazole, 10 mM sodium acetate, 10 mM glycylglycine, and 25 mM NaCl, with the pH adjusted between 3 and 10. The pH of each well was measured after TNS addition using a Tecan Spark multimode plate reader to read fluorescence (Ex 321 nm / Em 445 nm). Fluorescence data were fitted to the Henderson–Hasselbalch equation using GraphPad to determine the apparent pK=.

### SAXS

Measurements were performed at the SAXS BM29 beamline of the European Synchrotron Radiation Facility (ESRF, Grenoble, France). The beamline employed a Pilatus3 X 2M 2D detector. The sample-to-detector distance was set to 2.87 meters and the beam wavelength to 1.0 Å. The used flow cell consists of a 1.5 mm inner-diameter quartz capillary, with a 10 μm wall thickness. The samples in PBS buffer passed the SAXS cell at a flow rate of 0.2 mL/min, and such continuous flow was used to minimize radiation damage to the samples. The 2D data was integrated to yield 1D curves of intensity *I(q)*, in which *q* is the magnitude of the scattering vector *q = 4*π λ*^-1^ sin(*θ*/2)*, where λ is the wavelength of the beam and θ the scattering angle. The measured raw data is available through ESRF (<u>DOI: 10.15151/ESRF-ES1893861834</u>).

After background subtraction, the higher-*q* data (0.5 nm^-1^ < q < 4 nm^-1^) was modelled as a summation of Lorentzian peaks, using a power-law term to represent the background:

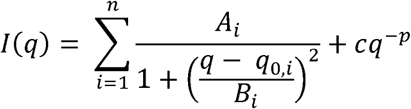

Where *n* is the total number of peaks, *A* is the amplitude (height) of each Lorentzian peak, *q_0_* is the center of the peak and *B* is the half-width at half-maximum (*hwhm*), *c* is a scale factor for the power-law background and *p* is its power. Two structure-independent parameters are derived from these fits: The characteristic distance *d =* 2π*/q_0_*, connected to the typical spacing of a structural feature, and the correlation length ξ = π*/B*, connected to the distance in which the structural ordering persists in space before becoming disoriented.

### Statistical analysis

Between-group comparisons were performed using one-way or two-way ANOVA with Tukey’s correction for multiple comparisons. QQ-plots were computed to investigate whether the data followed a Gaussian distribution. All statistical analyses were performed with GraphPad Prism 10.

## Acknowledgements

A.K. has received funding from the European Research Council (ERC) under the European Union’s Horizon Europe research and innovation programme (grant agreement No. 101118999, *Cat4CanCenter*, ERC-2023-SyG). As part of the COFUND project oLife, D. A. acknowledges funding from the European Union’s Horizon 2020 research and innovation program under grant agreement 847675. B.S and R.K. receive funding from ERA4Health and the Dutch Research Council (grant number era4healthcvd-112). I.K.V. acknowledges financial support of the European Research Council (ERC-2020-CoG 101001965) and the Dutch Ministry of Education, Culture and Science (Gravity program 024.005.020). Parts of figure 1-3 and 5 were created with BioRender.com using an academic lab subscription. The authors acknowledge the European Synchrotron Radiation Facility (ESRF) for provision of synchrotron radiation facilities under proposal number MX-2671 and would like to thank Dr. Stephanie Hutin for assistance and support in using beamline BM29. We thank Panagiota Papadopoulou for acquiring the initial cryo-TEM images.

## References

1. Blaser, H.-U. Chirality and its implications for the pharmaceutical industry. Rendiconti Lincei 24, 213–216 (2013).

2. Kallenborn, R. & Hühnerfuss, H. Enantioselective Toxic and Ecotoxic Effects of Drugs and Environmental Pollutants. in Chiral Environmental Pollutants 163–188 (Springer Berlin Heidelberg, Berlin, Heidelberg, 2001). doi:10.1007/978-3-662-06243-2_4.

3. Hutt,, Andrew J. CHIRALITY AND PHARMACOKINETICS: AN AREA OF NEGLECTED DIMENSIONALITY? Drug Metabol. Drug Interact. 22, 79–112 (2007).

4. Senkuttuvan, N. et al. The significance of chirality in contemporary drug discovery-a mini review. RSC Adv. 14, 33429–33448 (2024).

5. Spasov, A. A., Iezhitsa, I. N., Vassiliev, P. M., Ozerov, A. A. & Agarwal, R. Pharmacology of Drug Stereoisomers. vol. 76 (Springer Nature Singapore, Singapore, 2022).

6. Èižmáriková, R., Habala, L. & Valentová, J. General Anesthetics: Aspects of Chirality, Pharmacodynamics, and Pharmacokinetics. Pharmaceuticals 18, 250 (2025).

7. Kumari Rayala, V. V. S. P., Kandula, J. S. & P, R. Advances and challenges in the pharmacokinetics and bioanalysis of chiral drugs. Chirality 34, 1298–1310 (2022).

8. De, C. K. et al. The Overlooked Stereoisomers of the Ionizable Lipid ALC315. J. Am. Chem. Soc. 147, 28595–28600 (2025).

9. He, Z. et al. Unraveling the Role of Ionizable Lipid Isomerism in Modulating Lipid Nanoparticles for mRNA Delivery. J. Am. Chem. Soc. https://doi.org/10.1021/jacs.5c20438 (2026) doi:10.1021/jacs.5c20438.

10. Hatit, M. Z. C. et al. Nanoparticle stereochemistry-dependent endocytic processing improves in vivo mRNA delivery. Nat. Chem. 15, 508–515 (2023).

11. Da Silva Sanchez, A. J., et al. Substituting racemic ionizable lipids with stereopure ionizable lipids can increase mRNA delivery. Journal of Controlled Release 353, 270–277 (2023).

12. Dong, Y. et al. Lipopeptide nanoparticles for potent and selective siRNA delivery in rodents and nonhuman primates. Proc. Natl. Acad. Sci. U. S. A. 111, 3955–3960 (2014).

13. Zeng, Y., Escalona-Rayo, O., Knol, R., Kros, A. & Slütter, B. Lipid nanoparticle-based mRNA candidates elicit potent T cell responses. Biomater. Sci. 11, 964–974 (2022).

14. Koltover, I., Salditt, T., Rädler, J. O. & Safinya, C. R. An inverted hexagonal phase of cationic liposome-DNA complexes related to DNA release and delivery. Science (1979). 281, 78–81 (1998).

15. Chan, A. et al. Designing lipid nanoparticles using a transformer-based neural network. Nature Nanotechnology 2025 20:10 20, 1491–1501 (2025).

16. Kauffman, K. J. et al. Optimization of Lipid Nanoparticle Formulations for mRNA Delivery in Vivo with Fractional Factorial and Definitive Screening Designs. Nano Lett. 15, 7300–7306 (2015).

17. Dong, Y. et al. Lipopeptide nanoparticles for potent and selective siRNA delivery in rodents and nonhuman primates. Proceedings of the National Academy of Sciences 111, 3955–3960 (2014).

18. Sieber, S. et al. Zebrafish as a preclinical in vivo screening model for nanomedicines. Adv. Drug Deliv. Rev. 151–152, 152–168 (2019).

19. Ruchika, Sharma, A. & Saneja, A. Zebrafish as a powerful alternative model organism for preclinical investigation of nanomedicines. Drug Discov. Today 27, 1513–1522 (2022).

20. Arias-Alpizar, G., Bussmann, J. & Campbell, F. Zebrafish embryos as a predictive animal model to study nanoparticle behavior in vivo. Bio. Protoc. 11, 1–15 (2021).

21. Phillips, J. B. & Westerfield, M. Zebrafish models in translational research: Tipping the scales toward advancements in human health. DMM Disease Models and Mechanisms 7, 739–743 (2014).

22. Escalona-Rayo, O. et al. In vitro and in vivo evaluation of clinically-approved ionizable cationic lipids shows divergent results between mRNA transfection and vaccine efficacy. Biomedicine and Pharmacotherapy 165, 115065 (2023).

23. Pattipeiluhu, R. et al. Anionic Lipid Nanoparticles Preferentially Deliver mRNA to the Hepatic Reticuloendothelial System. Advanced Materials 34, 2201095 (2022).

24. Sago, C. D., Krupczak, B. R., Lokugamage, M. P., Gan, Z. & Dahlman, J. E. Cell Subtypes Within the Liver Microenvironment Differentially Interact with Lipid Nanoparticles. Cell. Mol. Bioeng. 12, 389–397 (2019).

25. Li, H. et al. Mesoscopic Structure of Lipid Nanoparticles Studied by Small-Angle X-Ray Scattering: A Spherical Core-Triple Shell Model Analysis. Membranes (Basel). 15, 153 (2025).

26. Yanez Arteta, M., et al. Successful reprogramming of cellular protein production through mRNA delivered by functionalized lipid nanoparticles. Proceedings of the National Academy of Sciences 115, (2018).

27. Sebastiani, F. et al. Apolipoprotein E Binding Drives Structural and Compositional Rearrangement of mRNA-Containing Lipid Nanoparticles. ACS Nano 15, 6709–6722 (2021).

28. Akinc, A. et al. Targeted Delivery of RNAi Therapeutics With Endogenous and Exogenous Ligand-Based Mechanisms. Molecular Therapy 18, 1357–1364 (2010).

29. Sahay, G. et al. Efficiency of siRNA delivery by lipid nanoparticles is limited by endocytic recycling. Nature Biotechnology 2013 31:7 31, 653–658 (2013).

30. Patel, S. et al. Naturally-occurring cholesterol analogues in lipid nanoparticles induce polymorphic shape and enhance intracellular delivery of mRNA. Nature Communications 2020 11:1 11, 983- (2020).

31. Tenzer, S. et al. Rapid formation of plasma protein corona critically affects nanoparticle pathophysiology. Nature Nanotechnology 2013 8:10 8, 772–781 (2013).

32. Voke, E. et al. Protein corona formed on lipid nanoparticles compromises delivery efficiency of mRNA cargo. Nature Communications 2025 16:1 16, 8699- (2025).

33. Li, C., Tan, X. F., Lim, T. K., Lin, Q. & Gong, Z. Comprehensive and quantitative proteomic analyses of zebrafish plasma reveals conserved protein profiles between genders and between zebrafish and human. Scientific Reports 2016 6:1 6, 24329- (2016).

34. Pattipeiluhu, R. et al. Liquid crystalline inverted lipid phases encapsulating siRNA enhance lipid nanoparticle mediated transfection. Nat. Commun. 15, 1–15 (2024).

35. Diez, M. et al. iCodon customizes gene expression based on the codon composition. Scientific Reports 2022 12:1 12, 12126- (2022).

36. Caysa, H. et al. A redshifted codon-optimized firefly luciferase is a sensitive reporter for bioluminescence imaging. Photochemical & Photobiological Sciences 8, 52–56 (2009).

37. Linares-Fernández, S. et al. Combining an optimized mRNA template with a double purification process allows strong expression of in vitro transcribed mRNA. Mol. Ther. Nucleic Acids 26, 945–956 (2021).

38. Zhuang, X. et al. mRNA Vaccines Encoding the HA Protein of Influenza A H1N1 Virus Delivered by Cationic Lipid Nanoparticles Induce Protective Immune Responses in Mice. Vaccines 2020, Vol. 8, 8, (2020).

39. Schindelin, J., et al. Fiji: An open-source platform for biological-image analysis. Nat. Methods 9, 676–682 (2012).

40. Aleström, P. et al. Zebrafish: Housing and husbandry recommendations. Lab. Anim. 54, 213–224 (2020).

41. Weinstein, B. M., Stemple, D. L., Driever, W. & Fishman, M. C. gridlock, a localized heritable patterning defect in the zebrafish. Nat. Med. 1, 1–6 (1995).

